# Molecular characterisation of neurons across animals identifies conserved gene expression

**DOI:** 10.1101/2025.06.04.657789

**Authors:** Carlos J. Rivera-Rivera, R. Feuda, D. Pisani

## Abstract

The evolution of the nervous system has been shrouded in controversy since the onset of the genomics era. A large part of this controversy stems from the lack of phylogenetic consensus for the main branches of the animal tree, where often animals with nervous systems do not form a monophyletic group. However, this question can be informed from other non-phylogenetic perspectives, such as comparative genomics. Here we ask how similar are genes differentially expressed in neurons across a representative set of eight animals with a nervous system and tally the presence or absence of their homologs in 10 other animals and two choanoflagellates. We show that proteins from 39 families are differentially expressed in all neurons, regardless of the phylogenetic placement of their lineage, and that the majority of these gene families are present in the (unicellular) closest relatives of animals, choanoflagellates. We found that the members of these 39 gene families are enriched in domains for ion transport and juxtacrine signalling, and that there is one gene family of zinc-dependent extracellular matrix-remodelling proteins which is only found in animals bearing a nervous system. Our results show that common genetic toolkits are in place for the function of nervous systems. We identify a large number of potential new genomic markers linked to the nervous system and hope they can complement ongoing research efforts to better understand this quintessentially animal system.

## Introduction

The nervous system, most commonly defined as a collection of neuronal cell types organised into a system via synaptic or anastomosed cell contacts [1], provided animals a discrete tissue system for integrating sensory information thus unlocking a free-living lifestyle in large multicellular organisms [2]. It can be argued that the nervous system is a hallmark of animal life and distinguishes the lineage in contrast to the other kingdoms of multicellular life.

Traditionally, it has been assumed that the nervous system is homologous across all animals. However, this hypothesis was recently challenged [3,4]. Uncertainty in our understanding of nervous system evolution follows from the emergence of incongruence at the root of the animal phylogeny [4–13] Traditionally, sponges (the Porifera), were considered the sister of all the other animals followed by the Placozoa (a small lineage of diploblastic marine organisms), and the Eumetazoa, traditionally identified as the lineage including all animals with a nervous system (Fig 1A, B, and E). Within Eumetazoa, two phyla; Cnidaria (corals and jellyfishes) and Ctenophora (comb jellies) were identified as sisters to each other in a lineage named Coelenterata. In turn, Coelenterata was identified as the sister to Bilateria (the lineage including all animals with bilateral body symmetry - acoel flatworms, arthropods and vertebrates - Fig. 1E). However, genomic data failed to corroborate this phylogeny, with some studies identifying the comb jellies as the sister of all the other animals (Fig. 1C and D) and others still identifying the sponges as the sister of all the other animals, but in a context where the comb jellies are the second lineage emerging along the animal tree (the tree in Fig. 1A and B). In these new scenarios, a single origin of the nervous system with no subsequent losses cannot be reconciled, suggesting more complex evolutionary histories where the nervous system might have evolved multiple times independently [14–17] or, if it evolved only once, it was then secondarily lost multiple times [4] (Fig. 1 summarises the possible scenarios), irrespective of the relative relationships of sponges and comb jellies.

**Figure 1.**
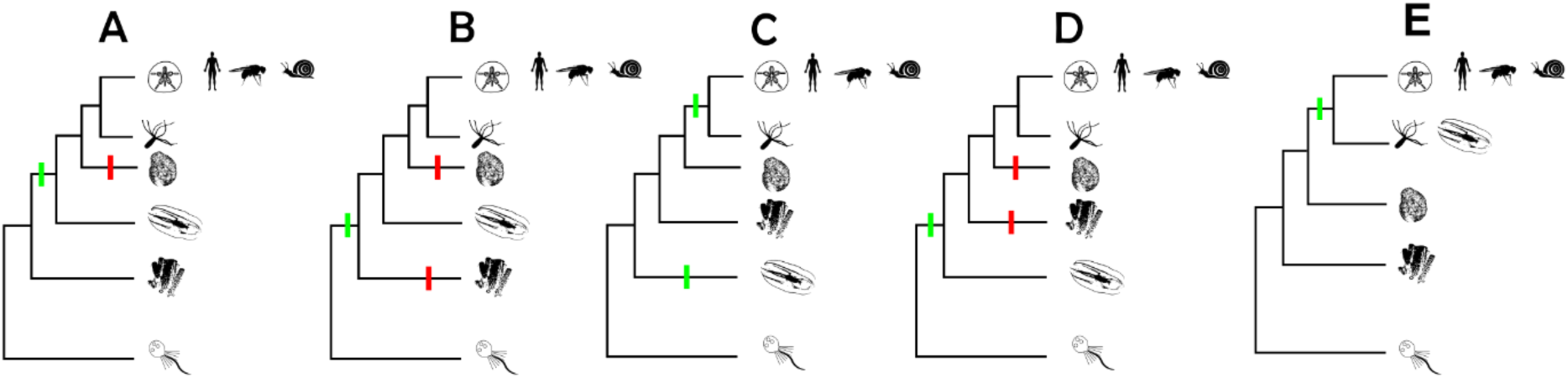
Proposed hypotheses for the emergence(s) of the nervous system(s). A - Assuming that sponges are the first metazoan lineage to branch out (Porifera-sister hypothesis), the nervous system emerged in the ancestor to all animals minus sponges and was lost in the ancestor to placozoans. B - Assuming Porifera-sister, the nervous system emerged in the ancestor to all animals and was independently lost in sponges and in placozoans. C - Assuming ctenophores are the first metazoan lineage to branch out (Ctenophora-sister hypothesis), at least two independent emergences occurred, at the ancestor of all ctenophores, and again in the ancestor to cnidarians and bilaterians. D - Again assuming Ctenophora-sister, nervous systems emerged in the ancestor to all animals and were subsequently lost in sponges and placozoans. E - Assuming the Coelenterata hypothesis (sponges as the first branching metazoan and cnidarians + ctenophores sister to bilaterians), the nervous system emerged in the ancestor to all animals with a nervous system.

Here we address the problem of the origin of the nervous system with reference to gene content. We investigate the evolutionary history of gene families with a special focus on gene orthogroups that are differentially expressed in the nervous system. We test how many such families are shared across different animal lineages and whether we could expect such numbers of shared families to be consistent with multiple origins of the nervous system. We used readily accessible whole-organism single-cell transcriptomes (scRNA-seq) for a variety of key taxa and the highly curated orthogroup database eggNOG [18]. We use scRNA-seq data from a ctenophore, three cnidarians, and four bilaterians to determine which gene orthogroups are differentially expressed in all nerve cells, how similar the orthogroups sets are to each other, and what their functional profiles are. We contrasted those results with the total coding sequence (CDS) data for 12 additional species spanning all main branches of animals and their closest relatives, the choanoflagellates, to determine at which taxonomic level each nervous system-expressed orthogroup emerged. Finally, we test whether the number of shared orthogroups is what we would expect if the nervous systems of key organisms evolved independently.

We found that irrespective of the relative relationships of ctenophores and sponges, all animals with a nervous system differentially express a core set of orthogroups in their neuronal cells. Pairwise tests showed that these sets are more similar to each other than expected by chance alone. We interpret these results to suggest that the nervous system most likely evolved only once, and was subsequently lost, in Placozoa (if sponges are the sister of all the other animals), or in sponges and Placozoa (if ctenophores are the sister of all the other animals). If it evolved twice independently, this underpinning similarity suggests that the same genes were independently co-opted in the evolving nervous systems. This interpretation is possible but less parsimonious.

## Materials and Methods

### Data acquisition and curation

We obtained post-processed unique molecular identifier (UMI) count tables from previously published scRNA-seq studies for eight metazoans with nervous systems: *Mnemiopsis leidyi* (Ctenophora) [19]*, Hydra vulgaris* (Cnidaria) [20]*, Nematostella vectensis* (Cnidaria) [21]*, Stylophora pistillata* (Cnidaria) [22]*, Crassostrea gigas* (Protostomia: Lophotrochozoa) [23]*, Drosophila melanogaster* (Protostomia: Ecdysozoa) [24]*, Mus musculus* (Vertebrata) [25,26], and *Danio rerio* (Vertebrata) [27]. For each, we also obtained the reference genomic CDS datasets used to annotate the scRNA-seq reads, as well as cell annotations where available (see Supplementary Table S1 for details of each dataset and references). The cell annotation for nervous system cells of *M. leidyi* was produced by [1], a different study from the one that initially produced the whole-organism scRNA-seq datasets. We also obtained full-organism CDS data for 12 other holozoan lineages: Choanoflagellata: *Monosiga brevicollis*, *Salpingoeca rosetta;* Porifera: *Spongilla lacustris, Amphimedon queenslandica, Ephydatia muelleri*; Ctenophora: *Pleurobrachia bacheii*; Placozoa: *Trichoplax adhaerens*; Platyhelminthes: *Schmidtea mediterranea, Schistosoma mansoni*; Echinodermata: *Strongylocentrotus purpuratus*; and Vertebrata: *Carcharadon carcharias, Xenopus tropicalis* (source dataset details in Supplementary Table S1). Combined with the eight species whose scRNA-seq data we analysed, we worked with a total of 20 species: two choanoflagellates and 18 metazoans. We analysed all 20 genome-level CDS datasets with eggnog-mapper v2.1.8 [28] and queried the eggNOG v5.0.2 [18] database with all the reference sequence sets using diamond v2.0.14 [29]. We then extracted, for each species, the list of orthogroups in the whole CDS set (‘totOGs’ from now on) based on their Opisthokonta-level annotation. Focusing on the Opisthokonta annotation allowed us to directly compare gene families across all species in our dataset. We also tallied the number of CDS without an eggNOG database hit, number of sequences without an Opisthokont orthogroup assignment, and the total number of orthogroups represented in each CDS dataset. These values allowed us to gauge the quality and completeness of our OG assignments (see Fig. 2 for a visual representation of the pipeline).

**Figure 2.**
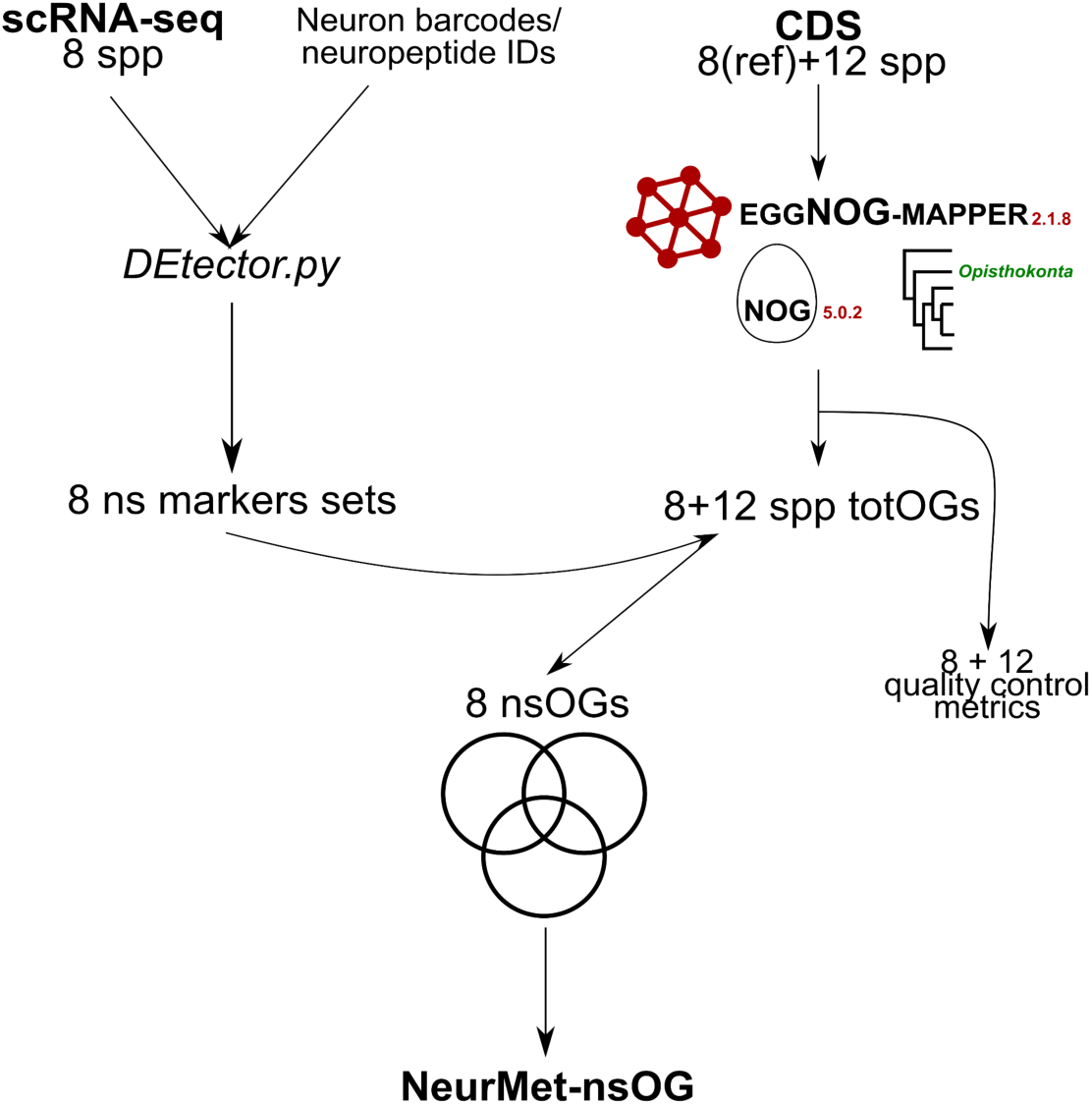
Schematic representation of the pipeline for obtaining the list of eggNOG orthogroups differentially expressed in nervous systems across animals. ScRNA-seq datasets for eight animal species were combined with the lists of neuronal cell barcodes or a list of neuropeptide markers (for *Crassostrea gigas*) in order to produce species-specific lists of genes differentially expressed in neurons. In parallel, the CDS reference datasets for all eight species plus 10 additional animals and two choanoflagellates were analysed with eggnog-mapper which provided the Opisthokonta-level orthogroup assignment for most CDS (totOGs). We gauged the quality and completeness of the eggnog-mapper results (see Fig. 3 below) to detect biases stemming from eggNOG 5.0.2’s database exhaustiveness. Next, each neuronal marker was queried in its corresponding CDS eggnog-mapper result, and the opisthokont-level orthogroup classification recorded. The resulting lists of nervous system orthogroups (nsOGs) were then compared with all sets against each other using in order to obtain the final list of 39 pan-metazoan orthogroups differentially expressed in neurons of the eight animals studied (‘NeurMet-nsOGs’).

### Pipeline for identifying nervous system cells

To detect differentially expressed genes in the scRNA-seq datasets we developed a bespoke script (available on https://github.com/carlosj-rr/scrnaseq_wrangle/DEtector.py). First, for each dataset and species, all genes which showed a stable expression pattern across cells (and as such were non-informative) were removed. These were identified as genes whose expression across cells showed a distribution statistically indistinguishable (using the Kolmogorov-Smirnov test) from a monotone dataset of the same size based on the median value. Next, we ranked differentially expressed genes and all UMI counts were recoded to ‘1’ for any cell in which the gene was expressed below the 5^th^ percentile of all cells (downregulation), ‘2’ for any cell in which the gene was expressed between the 5^th^ and 95^th^ percentiles of all cells (median expression), and ‘3’ for any cell in which the gene was expressed above the 95^th^ percentile of all cells (upregulation). We used a 5% cutoff to determine down- and upregulation because we wanted to focus on genes showing strong differential expression correlating with their neuronal cell type. These recoded tables (one per species) are available as Supplementary Tables S2-S9 in <u>data.bris</u>). We used two different approaches to identify nervous system genes from the table of differentially expressed genes. For all taxa but the mollusc *C. gigas* and the cnidarian *N. vectensis,* we extracted strongly differentially expressed genes in cell clusters identified as belonging to the nervous system in their original publication. For *N. vectensis*, the authors had identified a set of neural markers which we used to subselect the nervous system cells. For *C. gigas*, [23] did not identify neurons but identified 44 neuropeptide markers in their dataset. Based on that list of neuropeptides, we were able to identify nervous system cells (if they strongly differentially expressed more than 95% of the 44 neuropeptides).

### Detecting genes differentially expressed in nervous system cells

Next, we developed an algorithm and a metric to identify genes that are differentially expressed in neurons. First, we turned the recoded expression matrices S2 to S9 (see Supplementary Data) into Boolean tables by changing all ‘1’s and ‘3’s (down-/up-regulation) to ‘true’ and all ‘2’s (standard expression) to false. To detect genes with high differential expression in neuronal cells, we defined a metric that we named the proportional expression preference (PEP). This metric defines in what proportion of neuronal cells (‘foreground’ population) a gene is expressed in comparison to non-neuronal cells (the ‘background’ population).

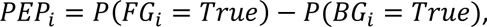

where 𝑃(𝐹𝐺*_i_* = 𝑇𝑟𝑢𝑒) represents the proportion of neuronal cells in which gene 𝑖 is differentially expressed, and 𝑃(𝐵𝐺*_i_* = 𝑇𝑟𝑢𝑒) the proportion of the non-neuronal cells in which the same gene is differentially expressed. This metric ranges from −1, if a gene is differentially expressed in all non-neuronal and zero neuronal cells, to 1, for a gene that is preferentially expressed in all neuronal cells and no non-neuronal cells.

Next, a permutation test was performed to define whether the observed PEP value for each gene was above what was expected purely by random expression noise: For each species, we counted the number of cells classified as nervous system cells. After that we randomly sampled the same number of cells from across the full dataset (nervous system cells included), 100 times. We then calculated the PEP values for each gene across the 100 random samples of cell populations. If the observed PEP value for a gene was above the 95^th^ percentile of the distribution of PEP values from random foreground/background cell combinations, the gene was identified as differentially expressed in nervous system cells. This resulted in the generation of species-specific lists of genes showing strong differential expression in nervous system cells.

To find common neuronal orthogroups among all species, we extracted the eggNOG opisthokont orthogroup annotation for each of the genes differentially expressed in neurons. In the eggNOG database, a gene family is equivalent to an orthogroup, as they encompass all orthologous and paralogous genes which descend from a common ancestor. Using the Opisthokonta-level orthogroup (OG) ID enabled us to compare OGs between all lineages in our dataset and allows for the results of our work to be compared to Fungi and other opisthokont clades that were not included in our analyses. This is important because, while Fungi and other opisthokonts do not have a nervous system, gene families with nervous system expression have deep evolutionary roots [30].

### Measuring similarities between sets of nervous system orthogroups and total genomic orthogroups at different taxonomic levels

We calculated the set intersection of total OG sets (totOGs) across all 20 CDS datasets. We determined, for our 20-species datasets, how many orthogroups were in common among vertebrates, protostomes, bilaterians, ctenophores, cnidarians, “coelenterates” (assuming their monophyly), eumetazoans, eumetazoans excluding the ctenophore, metazoans excluding the ctenophores, all metazoans (including ctenophores), and holozoans (metazoans plus choanoflagellates). We did the same analysis with the eight sets of OGs differentially expressed in neural cells (nsOGs), although the taxonomic sampling was limited to our eight species with scRNA-seq data.

Once we obtained a set of shared OGs across all metazoan nsOGs (NeurMet-nsOG), we defined at which point each neural cell OG emerged by cross-referencing with the taxonomic profile description in the online version of the eggNOG 5.0.2 database. We searched each NeurMet-nsOG’s taxonomic profile in the database’s website (http://eggnog5.embl.de/#/app/home; [18]), which shows the species in which a given OG is found, thus giving a good idea of when the OG may have emerged. We specifically asked which orthogroups were present in the last common ancestor of Fungi and animals, the last common ancestor of holozoans and animals, and within animals, the last common ancestor of all eumetazoans. Note that in the eggNOG 5.0.2 database Eumetazoa includes ctenophores by default. For all comparisons, we cross-referenced with our own datasets, to make sure our data corroborated or expanded those of the eggNOG database. This also enabled us to count the number of totOGs in common between all metazoans excluding ctenophores, a grouping that would emerge if the Ctenophora-sister hypothesis is true. Due to the way the eggNOG web interface is set up for the exploration of taxonomic profiles of gene families, we could not leverage the much larger taxonomic sample of that database for detecting totOGs which could have emerged in the ‘Metazoa minus ctenophora’ clade. In addition, we checked whether any of the NeurMet-nsOGs were present in the CDS sets of both choanoflagellates of this study.

### Functional profiling of NeurMet-nsOGs

To clarify the protein functional domains present in each NeurMet-nsOGs we used Pfam [31] and the InterPro database [32] for gene ontology (GO) annotation (where available).

### Quantifying difference between nsOG sets

We developed a permutation-based approach to test whether two nsOG sets were more similar to each other than expected by chance. Briefly, we extracted random subsamples of totOG of the same size as the nsOG set for each species and calculated the proportion of OGs found on both subsamples, relative to the size of the nsOG of a species of interest (100 permutations). We then tested whether the number of nsOGs shared between two species was significantly larger than that expected when randomly sampling OGs from these two species. To make sure the signal picked up in our analyses did not identify background similarity shared among cell types, we performed a validatory test using *M. leidyi*. For this test, we compared the OGs from the digestive cell cluster (dsOGs) of *M. leidyi* against the nsOGs of every other species in our dataset. We reasoned that if our results represented background similarity, rather than cell-specific homologous similarity, pairwise comparison between dsOGs and nsOGS should return results that are indistinguishable from those obtained when comparing nsOGs among each other.

## Results

### Nervous systems share 39 orthogroups

We identified the following numbers of nsOGs for the eight core species in our dataset: *C. gigas* (3,227), *D. melanogaster* (8,498), *D. rerio* (9,854), *H. vulgaris* (6,552), *M. leidyi* (1,125), *M. musculus* (7,023), *N. vectensis* (1,826), *S. pistillata* (3,889) (Table 1, neuron cell barcodes in Supplementary Table S10). Of these, a total of 39 were found in all eight species’ nsOG sets (see Supplementary Table S11 for each family’s name, based on mouse). We found that the most common biological process GO term among the 39 families was ‘transmembrane transport - GO:0055085’; for molecular function, ‘protein binding - GO:0005515’; and for cell compartment, ‘membrane - GO:0016020’ (full data in Supplementary Table S12). The most common protein domains in these 39 orthogroups include a large number of transmembrane proteins domains including protein, amino acid, carbohydrate, and ion transporters (mainly Ca^2+^, although in one case, an Na^+^ transporter, and in a different case H^+^ transporter) as well as GTPases, chaperones, cytoskeleton-associated proteins, kinases, and zinc fingers (Supplementary Table S12).

**Table 1.**
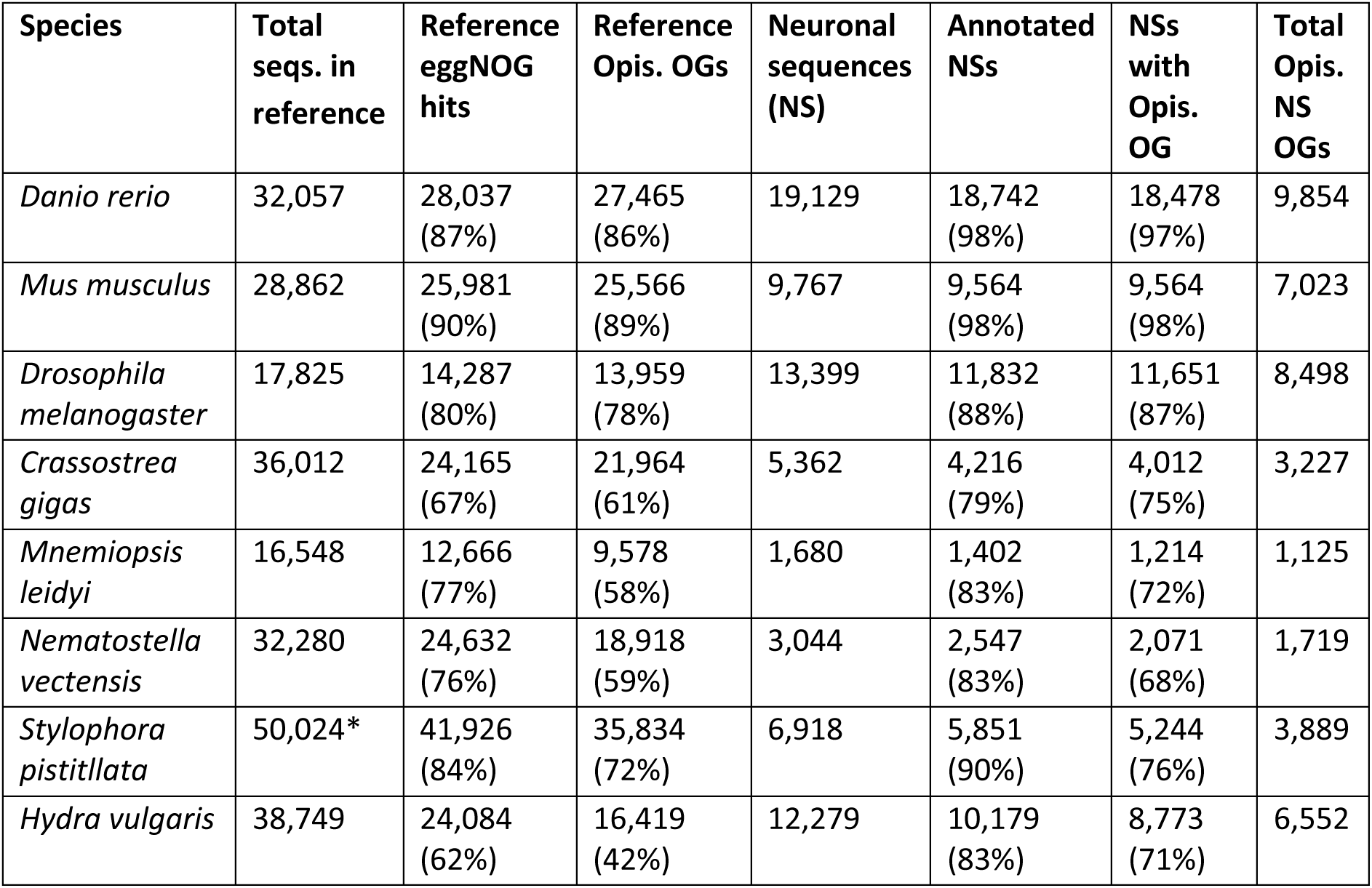
eggNOG search results overview. For all species that had scRNA-seq data analysed in this work, the total of sequences in the (CDS) reference dataset, how many of those were found in the eggNOG 5.0.2 database, how many of those had Opisthokonta-level orthogroup annotation. Then, how many genes were found to be differentially expressed in neurons, how many of those were found in the eggNOG 5.0.2 database, how may had an Opisthokonta-level orthogroup annotation, and finally, the total number of opisthokont orthogroups represented in the set of neuronally differentially-expressed sequences. The reference for *S. pistillata* included sequence data from several different sources and some sequences may have been repeated, which explains the large number of sequences in the reference. When tallying the total numbers of orthogroups (both for the CDS and neuronal datasets), repeated orthogroups were not allowed, which effectively removed repeats.

| Species | Total seqs. in reference | Reference eggNOG hits | Reference Opis. OGs | Neuronal sequences (NS) | Annotated NSs | NSs with Opis. OG | Total Opis. NS OGs |
| --- | --- | --- | --- | --- | --- | --- | --- |
| *Danio rerio* | 32,057 | 28,037 (87%) | 27,465 (86%) | 19,129 | 18,742 (98%) | 18,478 (97%) | 9,854 |
| *Mus musculus* | 28,862 | 25,981 (90%) | 25,566 (89%) | 9,767 | 9,564 (98%) | 9,564 (98%) | 7,023 |
| *Drosophila melanogaster* | 17,825 | 14,287 (80%) | 13,959 (78%) | 13,399 | 11,832 (88%) | 11,651 (87%) | 8,498 |
| *Crassostrea gigas* | 36,012 | 24,165 (67%) | 21,964 (61%) | 5,362 | 4,216 (79%) | 4,012 (75%) | 3,227 |
| *Mnemiopsis leidyi* | 16,548 | 12,666 (77%) | 9,578 (58%) | 1,680 | 1,402 (83%) | 1,214 (72%) | 1,125 |
| *Nematostella vectensis* | 32,280 | 24,632 (76%) | 18,918 (59%) | 3,044 | 2,547 (83%) | 2,071 (68%) | 1,719 |
| *Stylophora pistillata* | 50,024* | 41,926 (84%) | 35,834 (72%) | 6,918 | 5,851 (90%) | 5,244 (76%) | 3,889 |
| *Hydra vulgaris* | 38,749 | 24,084 (62%) | 16,419 (42%) | 12,279 | 10,179 (83%) | 8,773 (71%) | 6,552 |

We found that the species with the highest percentage of unique nsOGs was *D. melanogaster*, with 35.81% of its 8,498 nsOGs unique to its set (similar to the uniqueness of its totOGs set). The smallest set of unique nsOGs (7.20%) was that of *M. leidyi* (Fig. 3). The largest difference in uniqueness between the totOG and nsOG was found in *D. rerio*, for which 11.31% of its totOGs are lineage-specific, while 28.65% of its nsOGs are lineage-specific. *M. musculus* showed the smallest difference in number of unique OGs with 13.69% unique totOGs and 15.63% unique nsOGs (Fig. 2).

**Figure 3.**
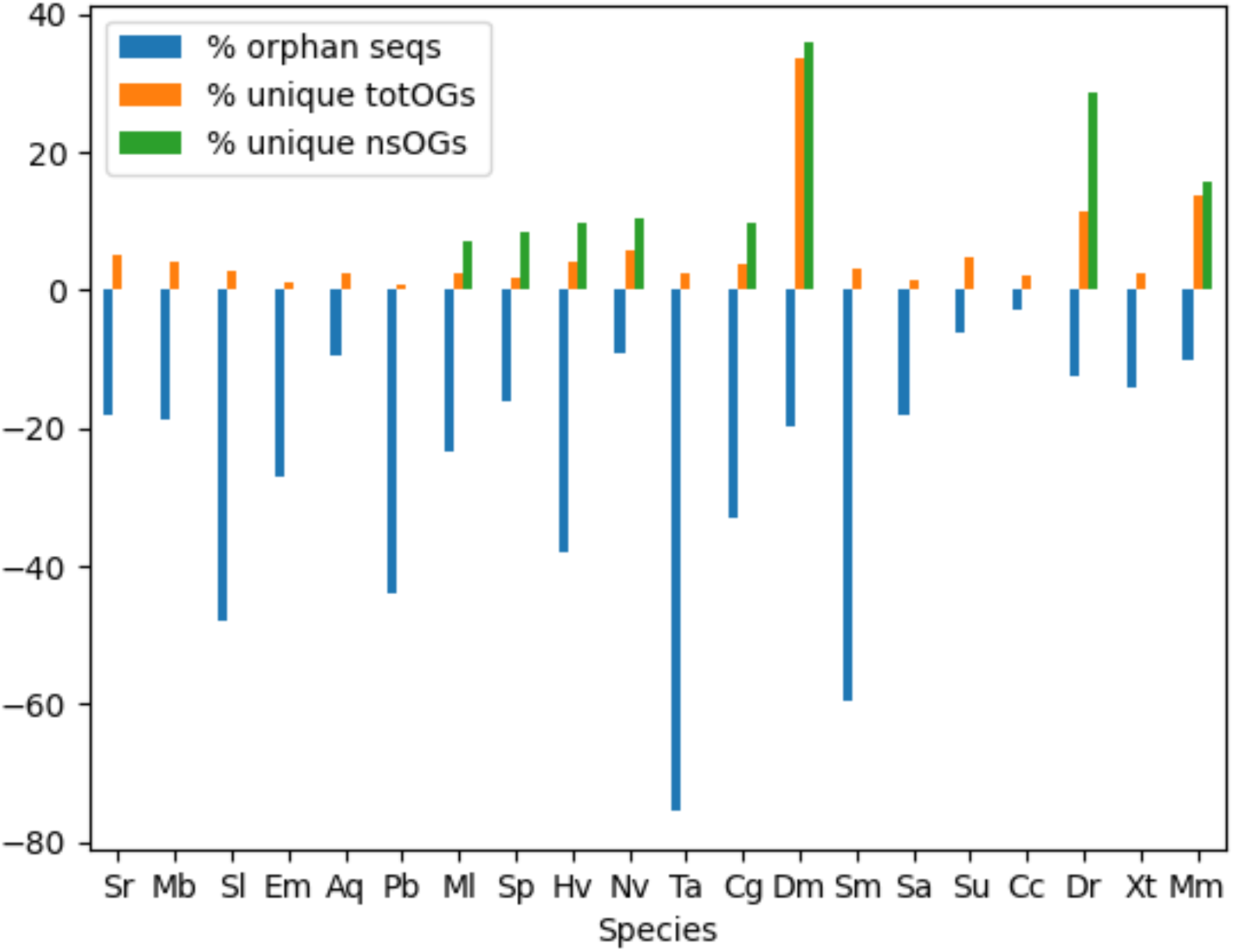
Genomic uniqueness for all species used in this study when compared to all other species in this study. Bars are shown for each species studied, and the colours represent the following: orange - percent of orthogroups in full CDS reference datasets which were unique to that species; green - percent of nervous system orthogroups which were unique to a particular species; and blue - percent of sequences (not orthogroups) which did not have a hit in the eggNOG 5.0 database. Species name codes are as follow: Sr - *Salpingoeca rosetta* (Choanoflagellata) Mb - *Monosiga brevicollis* (Choanoflagellata), Sl - *Spongilla lacustris* (Porifera), Em - *Ephydatia muelleri* (Porifera), Aq - *Amphimedon queenslandica* (Porifera), Pb - *Pleurobrachia bacheii* (Ctenophora), Ml - *Mnemiopsis leidyi* (Ctenophora), Sp - *Stylophora pistilliata* (Cnidaria), Hv - *Hydra vulgaris* (Cnidaria), Nv - *Nematostella vectensis* (Cnidaria), Ta - *Trichoplax adhaerens* (Placozoa), Cg - *Crassostrea gigas* (Mollusca - bilaterian), Dm - *Drosophila melanogaster* (Arthropoda – bilaterian), Sm - *Schimidtea mediterranea* (Platyhelminthes – bilaterian), Sa - *Schistosoma mansoni* (Platyhelminthes – bilaterian), Su – *Strongylocentrotus purpuratus* (Echinodermata – bilaterian) Cc - *Carcharadon carcharias* (Chordata – bilaterian), Dr - *Danio rerio* (Chordata – bilaterian), Xt - *Xenopus tropicalis* (Chordata – bilaterian), Mm - *Mus musculus* (Chordata – bilaterian).

### Orthogroups in common in CDS data studied

CDS sequences from the expanded dataset of 20 species with no hits in the eggNOG database ranged from 3.0% (the white shark) to 75.6% (the placozoan), (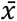 = 25.2%, 𝜎 = 18.6%, 𝑁 = 20, for average, standard deviation, and sample size, respectively – relative to all other species; see Supplementary Table S13a). The organism with the least number of totOGs is *T. adhaerens* (3,830), and the one with the most, *X. tropicalis* (12,514), (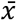 = 7,944, 𝜎 = 2,719.1, 𝑁 = 20). All 20 totOG sets (including two choanoflagellates and the placozoan) shared 437 orthogroups, whereas when constrained to only the species with a nervous system (14 in total, including all eight with scRNA-seq data, two flatworms, one more ctenophore, the white shark, and the purple sea urchin) this number is 1,258. In contrast, the totOG sets of the eight species with scRNA-seq data show 2,049 shared orthogroups. We also calculated the percent of the total set of orthogroups which was unique to the species when compared to all the other 19 totOG sets. The species with the highest percentage of unique totOGs was *D. melanogaster*, with 33.67% of its totOGs not found in any of the other 19 totOG sets. The species with the least percentage of unique totOG sets was *P. bacheii*, with 0.79% of its totOGs as lineage-specific. *D. melanogaster*’s percent of unique CDSs is also reflected in the percent of unique nsOGs, 35.81%. *M. leidyi*’s CDS reference had 2.50% of its totOGs as unique (see Fig. 3, and Supplementary Table S14a for fully detailed numbers). Note that these lineage-specific orthogroups are not innovations but rather retentions. We identified them from the eggNOG database of Opisthokont orthogroups, meaning that are likely present in other opisthokont lineages not studied here.

### Taxonomic distribution of global nsOGs

Of the 39 nsOGs found in common among all eight species of eumetazoans we studied (NeurMet-nsOGs), 84.6% were present in the choanoflagellate *S. rosetta* CDS data, and 76.9% in the choanoflagellate *M. brevicollis*. Sponges also showed transcriptoin of this set: *S. lacustris* (97.4%), *A. queenslandica* (89.7%), and *E. muelleri* (94.9%). Finally, in the placozoan *T. adhaerens*, which has a secondarily reduced body plan and no known nervous system [33] only 69.2% of the nsOGs were present in its CDS reference (Supplementary Table S13b). All deuterostomes in our dataset had 100% of the NeurMet-nsOGs in their CDS datasets (Fig. 4, Supplementary Table S13c).

**Figure 4.**
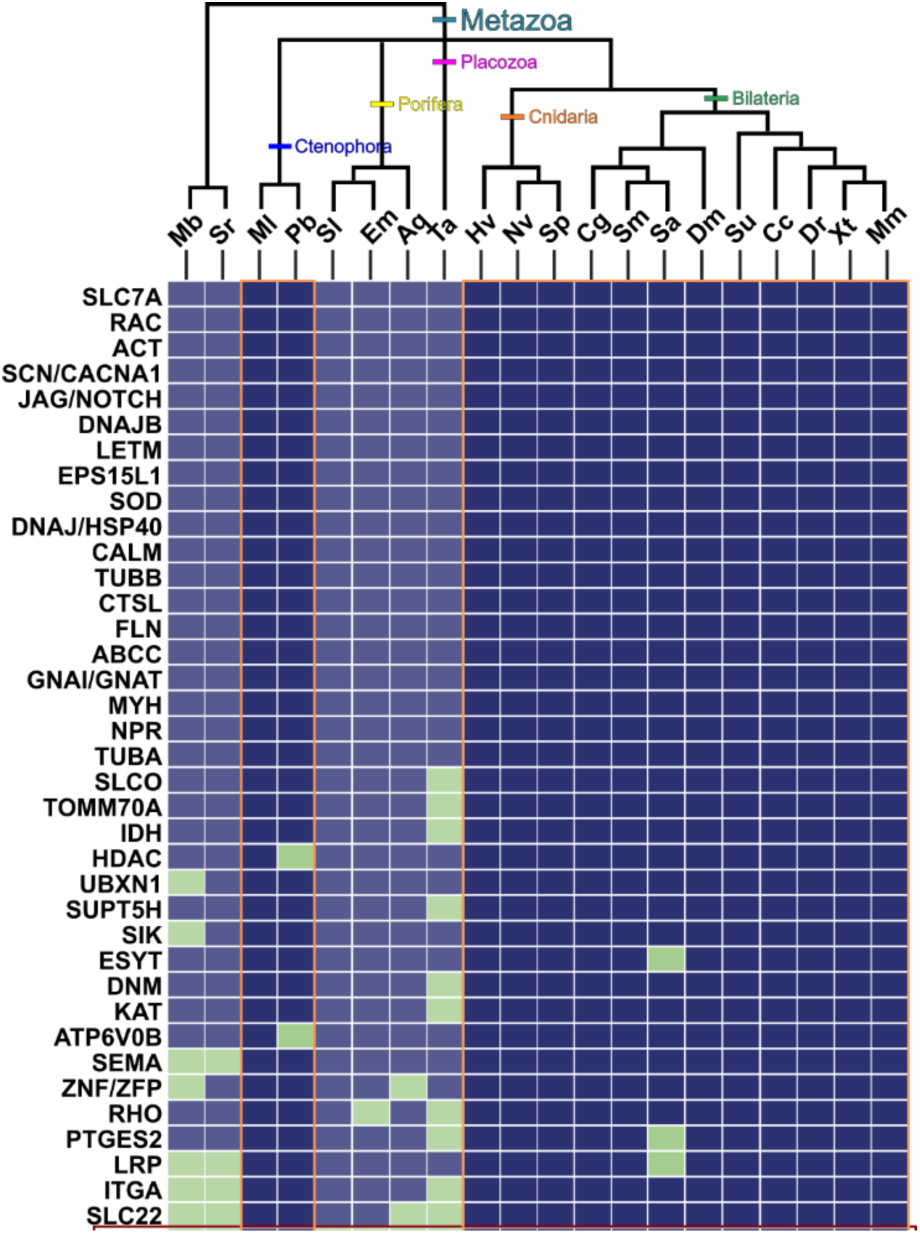
Presence/absence matrix of all 39 NeurMet-nsOGs in the CDS datasets of all the 20 species used in this study, using the following colour coding: Blue - orthogroup is present in the CDS dataset of that species; Green – orthogroup is absent in CDS dataset of that species. The species which are highlighted are species in our dataset which lack a nervous system and the OG highlighted (MMP) is one found only in organisms with a nervous system. OG names are based on mouse protein families.

According to the eggNOG taxonomic profiles, 29 of the 39 (74%) nsOGs shared by our core set of eight species were present in the common ancestor of animals and Fungi. Six (15%) emerged in the holozoan MRCA, three (7.5%) in the metazoan MRCA, and one (2.6%) in the ancestor to Eumetazoa (as defined in eggNOG database: i.e. Bilateria plus cnidaria and ctenophores, effectively, all animals with a nervous system). This eumetazoan specific nsOG is a member of the matrix metalloprotease (MMP – eggNOG code 38CRR@33154), a family of zinc-dependent extracellular matrix-remodelling proteins which belong to the metzincin protease superfamily [34] and we could confirm its absence in all three sponge CDS datasets as well as in *Trichoplax*’s (Fig. 4).

### NsOGs sets are more similar to each other than expected by chance

All pairwise comparisons among the nsOGs showed that the nsOGs of different species are more similar to each other than expected by chance, with all the observed values being larger than those from the random permutations (Supplementary Figure S1).

Our validatory test showed that there is usually greater similarity between nsOGs sets than between nsOGs and digestive system OGs (dsOGs). Indeed, for the cases of *D. melanogaster* and *H. vulgaris*, the number of OGs shared between the nsOG set and the dsOG set is random (Fig. 5). However, for the other cases it is not random. In the comparisons against the nsOGs of the remaining bilaterian taxa, the proportion of OGs shared between different nsOGs is higher than the proportion of OGs shared between dsOGs and nsOGs (Fig. 5). However, comparisons between the dsOGs of *M. leidyi* against the nsOGs of *N. vectensis* and *S. pistillata*, did not conform to the expected pattern, finding greater similarity than when comparing nsOGs among themselves. (Fig. 5).

**Figure 5.**
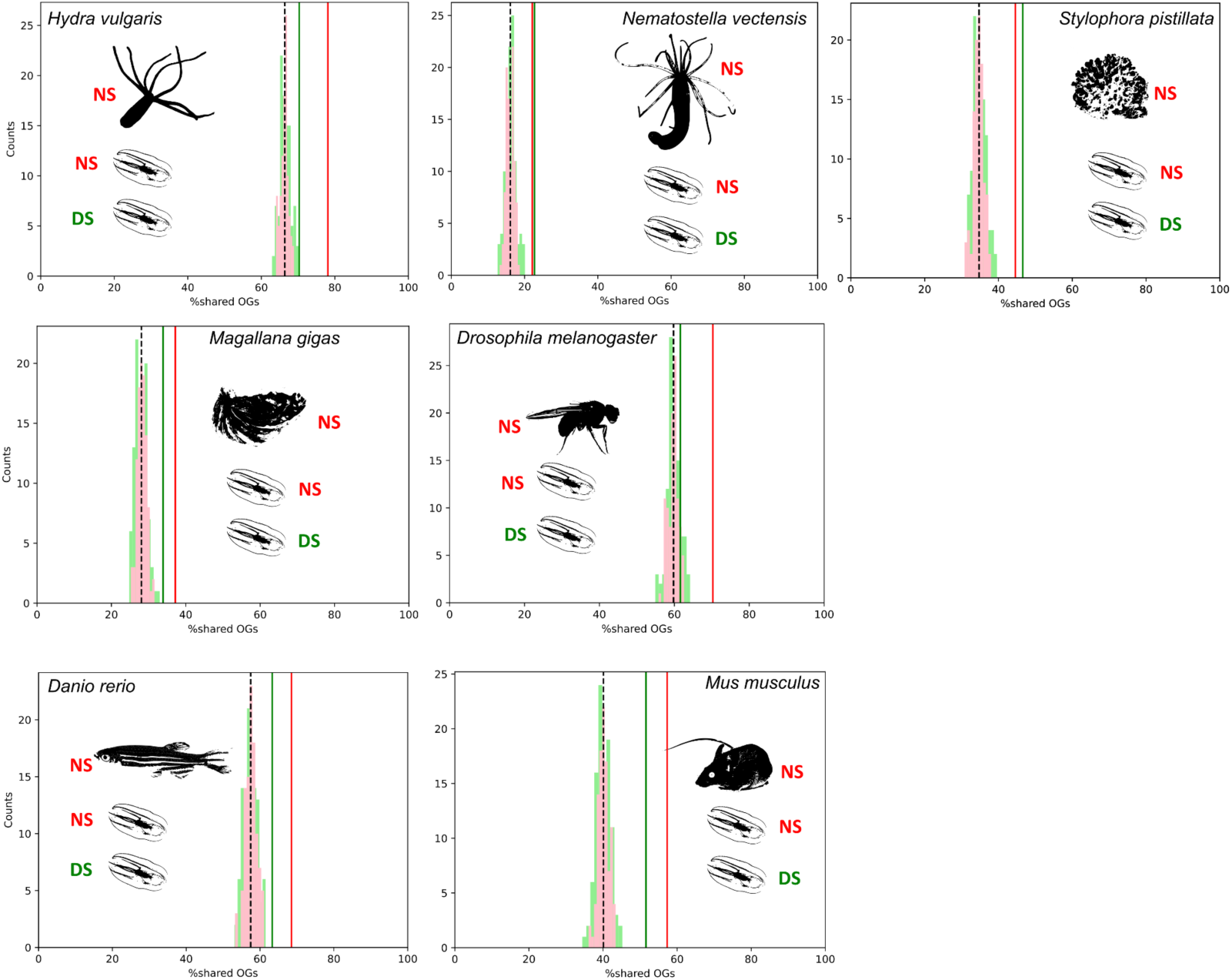
Overlap between the nervous and digestive system orthogroup sets of *M. leidyi* and the nervous system orthogroup sets of all other animals with a nervous system. Red distributions show the overlap between nervous systems expected by chance, and the red vertical line shows the observed overlap. Green distributions show the overlap expected by chance between the digestive system orthogroup set of *M. leidyi* and the second species’ nervous system orthogroup set, and the green vertical line is the observed overlap between both sets.

## Discussion

By combining scRNA-seq data and comparative genomics we show that 39 gene orthogroups have a common pattern of differential expression in neural cells. Furthermore, we showed that these orthogroups (four of which are Metazoan innovations) are enriched in protein domains that mediate cell-cell signalling as well as signalling within cells, but crucially, most of the orthogroups are present in choanoflagellates, the sister group to animals (84.6% and 76.9%, respectively). Finally, we showed that the set of nervous system orthogroups (nsOG) of the ctenophore *Mnemiopsis leidyi*, which has been previously hypothesised to have independently evolved its nervous system [3], is more similar to the set of the nsOGs of all other animals with a nervous system than is expected by chance. We compared these results with the total CDS data of 10 other animals and showed that most of these 39 nsOGs present in all nervous system-bearing animals are also found in animals without a nervous system (sponges and placozoans).

### Conservation/co-option of functional and developmental cues in animal nervous systems

Whether the nervous system emerged once at the root of the animal tree and was lost in sponges and placozoans, emerged independently in ctenophores and in the ancestor to cnidarians and bilaterians, or emerged once in an ancestor to ctenophores, cnidarians, and bilaterians after this lineage separated from sponges and placozoans, remains a fundamental but as of yet, unanswered question [4,11,13,35,36]. While part of this question can only be addressed phylogenetically (although ancestral syntenies were used by [4] - but see [13]), we can gain important clues about the origin of the nervous system through approaches that rely on the differential expression of gene families on neurons rather than on the relative placement of the main groups. We focus here on characterising the genomic component of neurons in general, with the intention of adding evidence to this question. Our results show that neurons share an enriched expression of domains for protein traffic and import/export, and ion channels. This does not come as a surprise, given the known role of neuropeptides and of paracrine signalling in whole-organism intercommunication [15,37], as well as the role of ion channels in creating actions potentials for neuronal firing [38]. It stands out that the neurons we studied here also share an enrichment in juxtacrine signalling domains. These domains are known to be essential mediators of cell differentiation during neurogenesis (Notch/Jagged, EGF-like, semaphorin), as well as crucial signals in axon guidance [39] and are present in all nsOGs. As more non-model organisms are studied using single-cell sequencing methods, this core set of genes can be revised and vetoed, and we look forward to a more comprehensive understanding of gene orthogroups involved in nervous system development and function.

### One nsOG is present in all nervous systems and absent from the CDS sets of animals lacking a nervous system

The taxonomic profile of one orthogroup, MMP (38CRR@33154) shows it to be unique to cnidarians, ctenophores, and bilaterians (both in the eggNOG database and in our own CDS data), suggesting a nervous system-specific retention that clusters all animals with a nervous system (Fig. 5). A second orthogroup, KANSL1 (38MCJ@33154) showed a similar pattern in the eggNOG database (unique to cnidarians, ctenophores, bilaterians, and placozoans), but we found it in two of our three sponge CDS datasets (*E. muelleri* and *S. lacustris*). The MMP-orthogroup sequences from *M. leidyi* are assigned by eggnog-mapper the KEGG pathway of ‘Parathyroid hormone synthesis, secretion, and action’ and the enriched Pfam domains for this OG across all animals were ‘Peptidase_M10’, ‘PG_binding_1’, ‘Hemopexin’, and ‘C1q’; domains related to peptide cleaving, bacterial cell wall degradation, haem binding, and activation of serum complement system, respectively (Supplementary Table S12).

### All nsOGs are more similar to each other than expected by chance

In pairwise comparisons, all nsOG sets were more similar to each other than expected by chance (Supplementary Figure S1). This is not surprising for the species which are hypothesised to have inherited the nervous system from a common ancestor, but it is interesting when considering the possibility that ctenophores independently evolved neurons and the nervous system. Assuming an independent emergence of the nervous system in ctenophores, one can envision that rudimentary neuronal components were already in place in the ancestor to all metazoans and thus were inherited by all clades (as discussed by [14]). These rudimentary neuronal components then evolved in parallel to form nervous systems which employ the same biochemical mechanisms. If this is the case, ctenophores, a group that in any case deserves much attention, will become a fundamental model organism for asking questions about the parallel evolution of complex systems in multicellular animals. On the other hand, if ctenophores inherited a fully integrated nervous system from an eumetazoan ancestor, they would still remain a fundamental group to study lineage-specific specialisation of nervous systems, as detailed structural and developmental studies have underlined [1,3,40]. However, these lines of inquiry will be contingent on a consensus on the phylogenetic placement of ctenophores in the animal tree of life.

### The evolution of a nervous system in an uncertain phylogenetic context

If we contextualise the results from this work into the most common phylogenetic hypotheses set forth for the early branching patterns of metazoans, we have the following possible scenarios. 1. Assuming the Ctenophora-sister hypothesis, the ancestor to all animals had a rudimentary signalling system that led to the parallel evolution of nervous systems with 39 orthogroups in common in ctenophores and then in all other animals, the ancestor of sponges lost its nervous system, and alongside it, the MMP orthogroup, and both the nervous system and the MMP orthogroup were again lost independently in the ancestor to placozoans (Fig. 6A). 2. Assuming the Ctenophora-sister hypothesis, the ancestor to all animals evolved a nervous system which already employed the 39 orthogroups found in common across all nervous systems, and this system was independently lost along with the MMP orthogroup in the ancestor to sponges and then to placozoans (Fig. 6B). 3. Assuming the Porifera-sister hypothesis, the ancestor to all non-poriferan metazoans developed a nervous system which used a new clade-specific orthogroup, MMP, which the ancestor to placozoans subsequently lost alongside its nervous system (Fig. 6C). 4. Assuming the Porifera-sister hypothesis, the ancestor to all animals developed a rudimentary signalling system which was lost in the ancestor to sponges, but the ancestor to all animals except sponges retained along with the MMP orthogroup, and subsequently evolved into which the ancestor to placozoans lost in its morphological reduction (Fig. 6D). 5. Assuming the Coelenterata hypothesis, the ancestor to all animals with a nervous system evolved an integrated nervous system, which included the MMP orthogroup (Fig. 6E). A recent study in Placozoan peptidergic neurons showed support for the hypothesis that the earliest instance of a nervous system was a secretory network of cells, which lends support to the hypotheses presented in Fig. 6A and 6C [33] (Najle et al. 2023).

**Figure 6.**
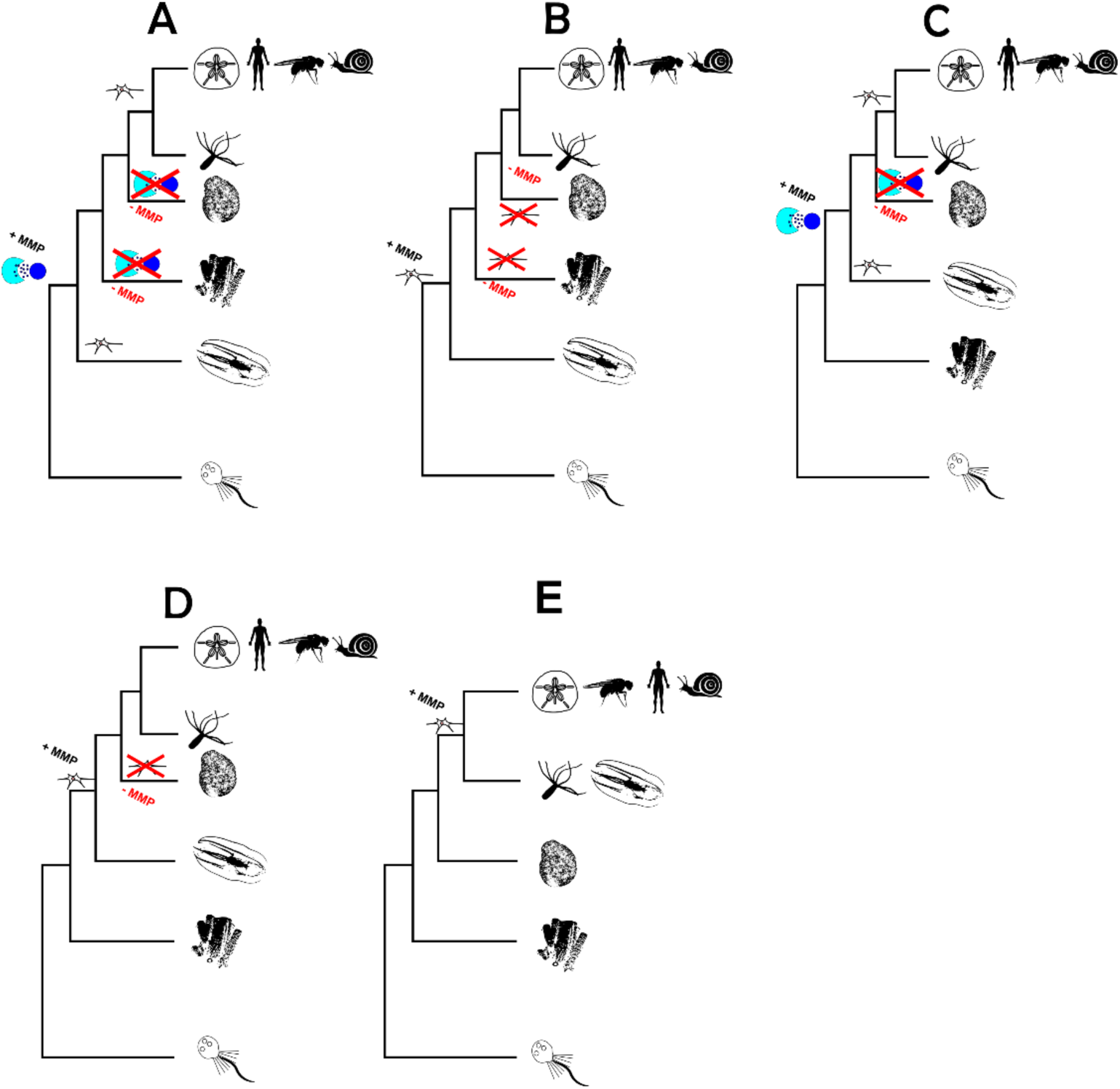
The five phylogenetic hypotheses discussed in this paper with the data from this study included in them. The hypotheses are as follows: A. Assuming Ctenophora-sister, a rudimentary paracrine signalling nervous proto-system which included the MMP OG was present in the ancestor to all animals. This nervous proto-system evolved into the nervous system we observe in ctenophores nowadays, and on its sister branch, it was lost independently (at least as a purely nervous signalling system), alongside the MMP OG, two times – in Porifera and in Placozoa. Finally, in the ancestor to Bilateria and Cnidaria, the nervous proto-system evolved into the nervous system we see in these groups. B. Assuming Ctenophora-sister, the ancestor to all animals already had a fully integrated nervous system which included the MMP OG, and this system, along with the MMP gene family was lost independently in Porifera and Placozoa. C. Assuming Porifera-sister, a nervous proto-system and the MMP OG were present in the ancestor to all non-poriferan animals, which then independently evolved a fully-integrated nervous system in Ctenophora and Bilateria + Cnidaria, and Placozoans lost the MMP OG as well as the nervous proto-system. D. Assuming Porifera-sister, the ancestor to all non-sponge animals evolved a fully-integrated nervous system that used the MMP OG in its development/operation, and this system and the OG were lost in Placozoa. E. Assuming the Coelenterata grouping, the ancestor of Bilateria + Coelenterata evolved a fully integrated nervous system which used the MMP OG in its development and/or operation.

Here we completed the first detailed description of the orthogroup composition of metazoan nervous systems, using scRNA-seq data from representative lineages of eight main metazoan branches. We showed how, regardless of phylogeny, animal nervous systems share important components of their biochemical machinery both for development and function, and we show that a single orthogroup which is differentially expressed in all nervous systems is present only in the genomes of animals with a nervous system and absent otherwise. This work represents an essential first step into reconstructing the regulatory networks conserved/co-opted in nervous systems across animals, which can be a key non-phylogenomic source of information to clarify the contentious evolutionary path of this quintessential animal system.

## Data access

All scRNA-seq and CDS data were obtained from public databases (see Supplementary Table S1 for sources). All supplementary figures, tables and data are available at the University of Bristol data repository, <u>data.bris</u>, at ok, the dataset has been published, although it may take some time for it to be seen here: https://doi.org/10.5523/bris.3n4sygf5c6fii2exgd5zgcyorv. All code to replicate the analysis, with detailed explanations on how each function works is available in CJRR’s GitHub repository: github.com/carlosj-rr/scrnaseq_wrangle.

## Acknowledgements

This work was funded by a Marie Skłodowska-Curie Action to CJRR (project code RipGEESE), RF is supported by the Royal Society (UF160226 and URF/R/221011) and DP and RF are supported by a Leverhulme Trust Grant RPG 2024-030. We thank all the members of the Bristol Palaeobiology group for their comments, observations, and encouragement during this work. All the high-performance computational analyses were carried out using the facilities of the Advanced Computing Research Centre, University of Bristol - http://www.bristol.ac.uk/acrc/.

